# Four temperate phages of the coral pathogen *Vibrio coralliilyticus* represent two novel genera

**DOI:** 10.64898/2026.09.21.753358

**Authors:** Jacob A. Albee, Nathan Fowler, Julianne H. Grose, Jing Yan, Colby Weeks

## Abstract

*Vibrio coralliilyticus* is an emerging coral pathogen associated with tissue-loss diseases in reef-building corals and with infections in other marine organisms, including oyster larvae. This study describes four novel *Vibrio* phages, Satoshi, Kolob, Ushijima, and Kane, isolated near Lā‘ie, Hawai‘i, on *V. coralliilyticus* strain OCN008, an etiological agent of *Montipora* white syndrome. These are the first isolated *V. coralliilyticus* phages described as temperate and the first from O‘ahu. Genome organization and taxonomic relationships were examined using whole-genome sequencing and comparative genomics, while representative morphologies were examined by transmission electron microscopy. Comparative analyses place all four phages within the class *Caudoviricetes* but outside any previously described orders, families, or genera. Satoshi, Kolob, and Ushijima cluster with the previously described *V. coralliilyticus* phage CKB-S2 in one genus-level lineage spanning Hawai‘i and Okinawa (proposed genus *Mokushimavirus*), while Kane, which shares at most approximately 1% of its sequence with any viral genome in public databases, represents a distinct lineage (proposed genus *Laievirus*). All four phages encode homologs of virulence factors that may contribute to host pathogenicity during lysogeny.

**IMPORTANCE:** *Vibrio coralliilyticus* threatens reef-building corals and oyster larvae, and its phages have been explored for phage therapy. Yet none of its more than two dozen phages isolated to date have been described as temperate. Temperate phages can introduce virulence genes into their hosts, and genomic analyses already predict toxin-like genes on prophages in some strains. This study reports the first isolated phages of this pathogen described as temperate, representing two previously undescribed genera, one with no known close relatives. These phages also carry homologs of virulence-associated genes, including some linked to antimicrobial peptide resistance, which could help their host withstand coral defenses if expressed during lysogeny. They provide both taxonomic reference points and the first opportunity to test experimentally how lysogeny may shape coral disease.

## INTRODUCTION

Global coral cover has declined by nearly 50% since the mid-20th century [1], with coral diseases accounting for an estimated *∼*30% of losses, alongside other anthropogenic and environmental stressors [2]. Among coral pathogens, *Vibrio* species are particularly prominent [3], including *Vibrio coralliilyticus* (*V. coralliilyticus*), an emerging pathogen of reef-building and soft corals, oyster larvae, and other bivalve larvae [4, 5]. *V. coralliilyticus* has been implicated in coral bleaching, tissue loss, and disease outbreaks across reef ecosystems worldwide [5], and has been described as the “best-known bacterial pathogen of corals” [6].

*V. coralliilyticus* was first described in the early 2000s following its isolation from diseased corals off the coast of Zanzibar, East Africa [5, 7]. Since its initial characterization, strains of *V. coralliilyticus* have been isolated across a broad geographic range, including the Indo-Pacific region [8], the Caribbean [9], and the Mediterranean Sea [10]. In the Hawaiian Islands, the indigenous *V. coralliilyticus* strain OCN008 has been established as an etiological agent of acute *Montipora* white syndrome—a rapidly progressing tissue-loss disease affecting the major reef-building coral *Montipora capitata* (*M. capitata*)—and is associated with substantial coral mortality in Kāne‘ohe Bay, O‘ahu [11].

*V. coralliilyticus* is also subject to ecological pressures exerted by bacteriophages, which influence the abundance, diversity, and evolution of bacterial populations [12]. Virulent phages drive bacterial evolution by killing infected cells, whereas temperate phages can follow either a lytic or a lysogenic cycle, the latter involving integration into the host genome as a prophage [12]. Prophage carriage can influence bacterial fitness, genomic diversification, susceptibility to subsequent phage infection, and pathogenicity [13].

*Vibrio* phage YB2, recovered from the Red Sea, was the first phage reported to infect *V. coralliilyticus* [14]. Since its discovery in 2007, at least 24 additional *V. coralliilyticus* phages have been described, from locations as diverse as Australia [5], Japan [15], South Korea [16–18], Vietnam [19], and, most recently, from a shellfish hatchery in Kailua-Kona, Hawai‘i Island [20].

Phages infecting *V. coralliilyticus* frequently exhibit a high degree of host specificity, infecting only a narrow subset of bacterial strains [20]. Despite the clear pathogenic significance of *V. coralliilyticus* OCN008, only one previously described phage (vB_VcorM-GR28A) has been documented as infecting this strain [20], and no *V. coralliilyticus* phages have been isolated from O‘ahu, where OCN008 was discovered [21].

Studies of *V. coralliilyticus* phages have focused on those with virulent lifestyles, largely as potential biocontrol agents [5, 14, 15, 18–20], reflecting the broader preference for lytic phages in phage therapy research [22], whereas the ecological consequences of lysogeny remain comparatively underexplored [23]. Prophages are widespread in vibrios (5,674 prophage-like elements identified across 1,874 genomes [24]), and can transfer virulence-associated genes that affect host pathogenicity [25]. In *V. coralliilyticus*, temperate phages have been inferred only from prophages predicted in sequenced bacteria, some encoding toxin-like genes [26], and, to date, no isolated phage has been described as temperate. Without isolates, a predicted prophage remains a candidate: its sequence alone cannot establish whether it is functional, active, or able to infect other cells, and many prophages are defective [27, 28]. By contrast, isolated phages are infectious and can be studied directly, providing the experimental material needed to test what genomic predictions can only suggest [26].

This study describes four novel phages (*Vibrio* phages Satoshi, Kolob, Ushijima, and Kane), isolated near Lā‘ie, Hawai‘i, that infect *V. coralliilyticus* OCN008 and are predicted to be temperate. These are the first cultured *V. coralliilyticus* phages to be described as such and the first reported from the island of O‘ahu. This study broadens our understanding of the pathogen’s viral diversity, provides the basis for proposing two new genera within the class *Caudoviricetes* (*Mokushimavirus* and *Laievirus*), and identifies candidate virulence factors that may influence the disease potential of this major coral pathogen.

## RESULTS

### Morphological Characterization of Phages

Four phages infecting *V. coralliilyticus* strain OCN008, denoted as *Vibrio* phages Satoshi, Kolob, Ushijima, and Kane, were isolated from coastal seawater collected at Temple Beach in Lā‘ie, Hawai‘i. All four *Vibrio* phages produced extremely turbid, barely visible plaques no more than 1.0 mm in diameter.

Negative stain transmission electron microscopy (TEM) was performed on *Vibrio* phages Ushijima and Kane, which represent the two genus-level groups defined by comparative genomics below: the Satoshi-Kolob-Ushijima group (proposed genus *Mokushimavirus*) and Kane (proposed genus *Laievirus*).

TEM revealed that *Vibrio* phage Ushijima (Fig. **1**A) exhibited a podovirus-like morphology, with a short tail and capsid structure that appeared icosahedrally symmetric.

**FIG 1.**
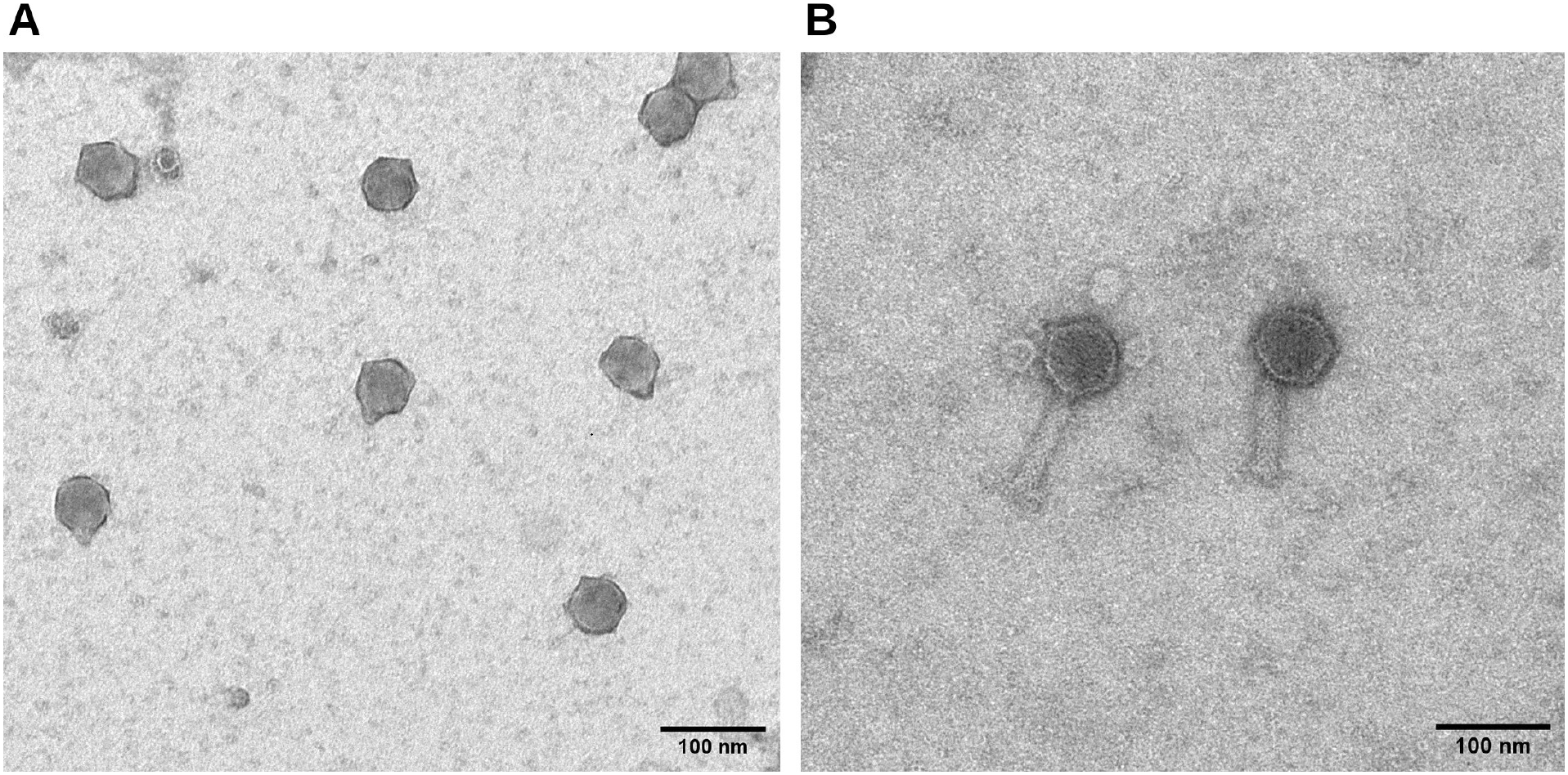
Transmission electron micrographs of *Vibrio* phages (A) Ushijima and (B) Kane.

In contrast, *Vibrio* phage Kane (Fig. **1**B) exhibited a myovirus-like morphology, with an icosahedrally symmetric capsid and a long, contractile tail with a visible sheath.

Dimensions of both phages are in Table **1**.

**TABLE 1.** Genome and virion features of *Vibrio* phages Satoshi, Kolob, Ushijima, and Kane. CDSs, predicted coding DNA sequences. Capsid and tail dimensions are mean *±* standard deviation from negative-stain TEM; Kane’s tail length excludes tail fibers. Satoshi and Kolob were not imaged, and a dash indicates a value not reported. Satoshi, Kolob, and Ushijima encode the same 21 named products.

| **Feature** | **Satoshi** | **Kolob** | **Ushijima** | **Kane** |
|---|---|---|---|---|
| Genome length (bp) | 34,888 | 34,556 | 34,423 | 55,994 |
| G+C content (%) | 45.2 | 45.3 | 45.6 | 44.7 |
| Predicted CDSs | 47 | 47 | 49 | 84 |
| on the predominant strand | 47 | 47 | 49 | 73 |
| on the opposite strand | 0 | 0 | 0 | 11 |
| Hypothetical proteins (%) | 55.3 | 55.3 | 57.1 | 53.6 |
| Proteins with predicted functions | 21 | 21 | 21 | 39 |
| Capsid diameter (nm) | — | — | 55.0 ± 2.2 | 63.5 ± 1.6 |
| Tail length (nm) | — | — | 12.2 ± 1.4 | 96.2 ± 1.5 |
| Tail width (nm) | — | — | — | 20.6 ± 1.2 |
| Particles measured (*n*) | — | — | 10 | 8 |

### Genomic Characterization and Taxonomic Placement

The genomes of *Vibrio* phages Satoshi, Kolob, Ushijima, and Kane are linear, circularly permuted double-stranded DNA (Table **1**). The G+C contents of the four phages (44.7–45.6%) were similar to that of the host strain *V. coralliilyticus* OCN008 (45.7%) [29].

BLASTn queries against the NCBI Virus database identified *Vibrio* phage CKB-S2 as the closest known relative of Satoshi, Kolob, and Ushijima. CKB-S2 (AP014888.1) is a phage infecting *V. coralliilyticus* strain P1 that was originally isolated from Okinawa, Japan [15].

VIRIDIC analysis of pairwise intergenomic nucleotide similarity (Fig. **2**) found 80.7–83.9% similarity between CKB-S2 and each Hawaiian phage and 77.7–87.2% among the Hawaiian phages, with Satoshi and Kolob the most similar.

**FIG 2.**
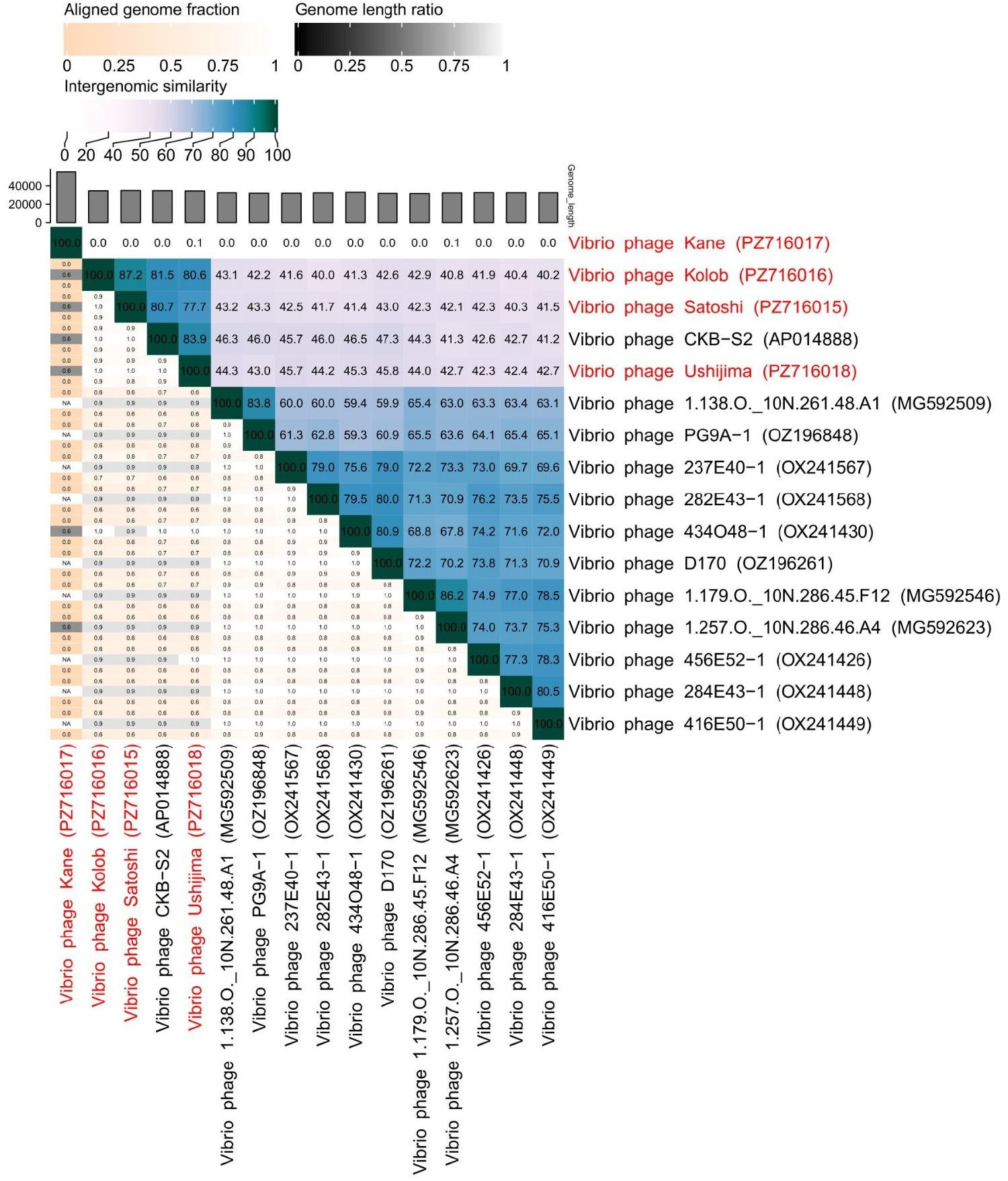
Pairwise intergenomic similarities (VIRIDIC) among the four novel phages (red) and selected reference *Vibrio* phages. Upper right: percent similarity. Lower left, top to bottom in each cell: aligned fraction of the row genome, genome length ratio, and aligned fraction of the column genome. Bars show genome length (bp).

Two independent approaches, VIRIDIC (nucleotide similarity) and vConTACT3 (gene-sharing networks) each assigned Satoshi, Kolob, Ushijima, and CKB-S2 to the same genus-level cluster. vConTACT3 assigned all four viruses to the class *Caudoviricetes* but did not place the cluster within any currently recognized order, family, subfamily, or genus.

Beyond CKB-S2, the closest available relatives of Satoshi, Kolob, and Ushijima in the NCBI Virus database included phages isolated on *Vibrio chagasii* and on *Vibrio crassostreae* during separate 2017 and 2021 surveys at an oyster farm near Brest, France, as well as phages isolated on *Vibrio lentus* and on *Vibrio splendidus* from coastal seawater at Canoe Cove, Nahant, Massachusetts [30–32].

Genome-wide proteomic analysis using ViPTree (Fig. **3**) placed Satoshi and Kolob as sister taxa, consistent with their high intergenomic similarity. The Satoshi-Kolob pair formed a larger clade with Ushijima, while CKB-S2 was sister to the clade comprising the three Hawaiian phages. Together, these four phages formed their own cluster, separate from the remaining reference phages. Maximum-likelihood phylogenetic analysis of terminase large subunit (TerL) amino acid sequences performed in MEGA12 (Fig. **4**) reproduced the same nested topology.

**FIG 3.**
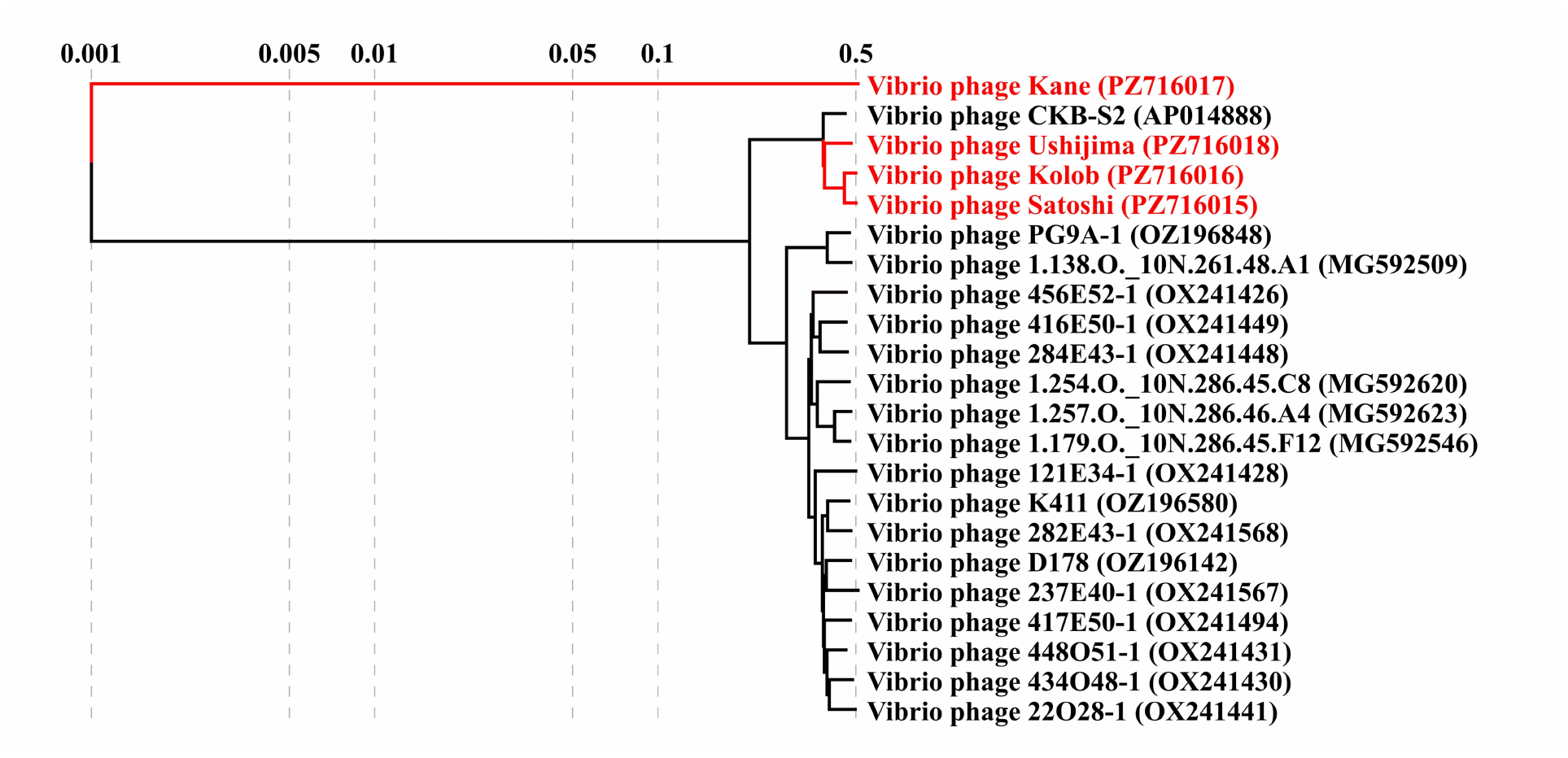
Proteomic tree of the four novel phages (red) and selected reference *Vibrio* phages (black), generated with ViPTree. Branch lengths are genomic distances on a logarithmic scale.

**FIG 4.**
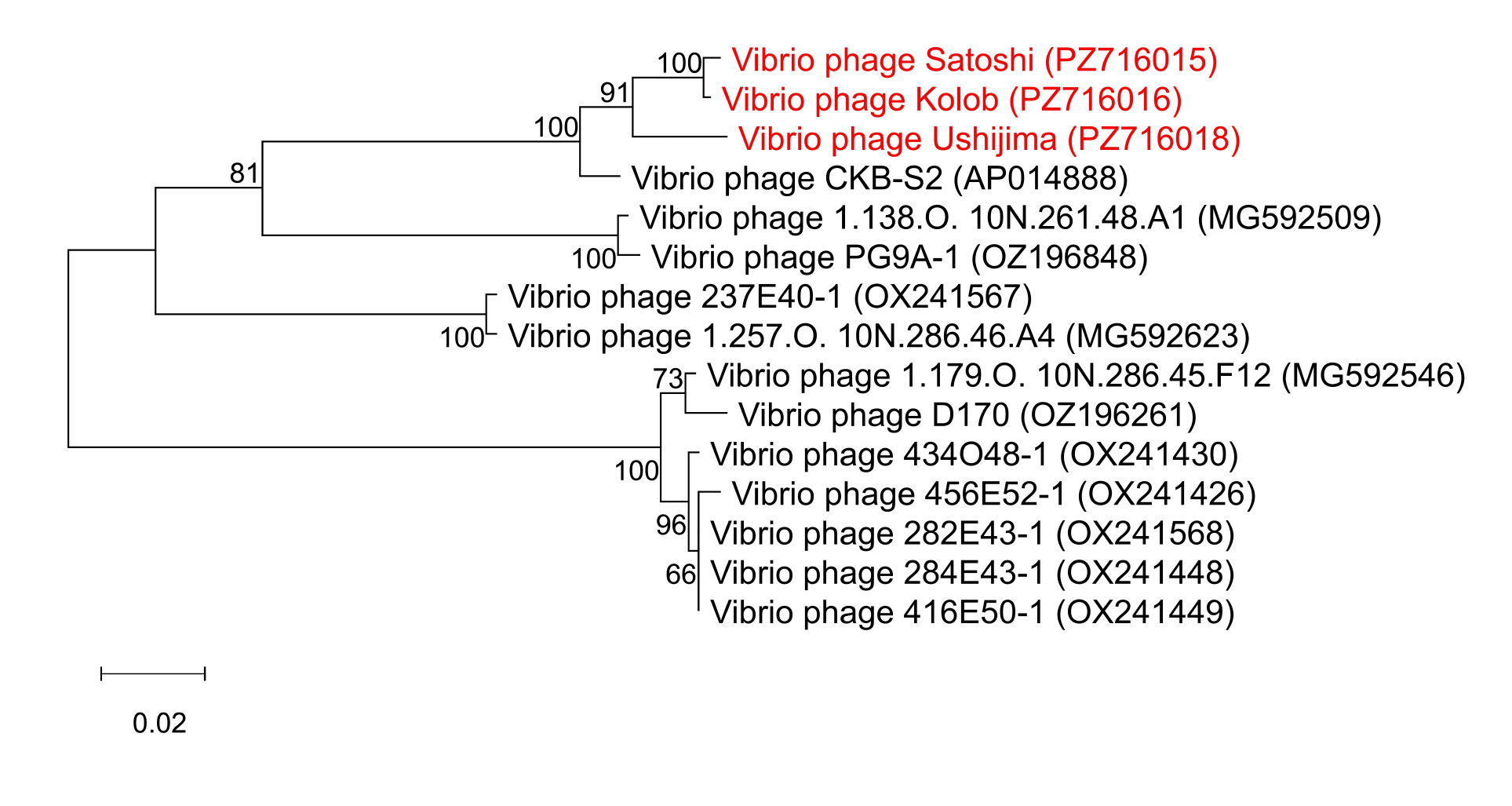
Maximum-likelihood phylogeny of terminase large subunit (TerL) amino acid sequences from *Vibrio* phages Satoshi, Kolob, and Ushijima (red) and relatives. Numbers at nodes are bootstrap percentages (1,000 replicates); the scale bar indicates amino acid substitutions per site. Kane was excluded because no close TerL relatives were found.

Pairwise intergenomic similarities among *Vibrio* phages Satoshi, Kolob, Ushijima, and CKB-S2 (Fig. **2**) fell below the 95% species-level threshold but above the 70% genus-level threshold commonly used for bacteriophage taxonomy [33], supporting their classification as distinct species within a novel genus. A formal taxonomic proposal has been submitted to the International Committee on Taxonomy of Viruses (ICTV) to establish this genus, named *Mokushimavirus*, and its four associated species, *Mokushimavirus satoshi*, *Mokushimavirus kolob*, *Mokushimavirus ushijima*, and *Mokushimavirus ckbs2*. The name *Mokushimavirus* combines the Hawaiian *moku* and Japanese *shima*, both meaning “island,” in reference to the Hawaiian and Japanese origins of its four member species.

In contrast to the *Mokushimavirus* members, BLASTn searches identified no close relatives of phage Kane. The highest-scoring matches, which consisted predominantly of *Vibrio* phages, produced no more than approximately 1% query coverage. VIRIDIC analysis of these matches revealed extremely low intergenomic similarities, the highest being only 0.8%, observed between Kane and *Vibrio* phage vB_ValR_NF (MN812722), a *Vibrio alginolyticus* phage from coastal waters near Qingdao, China [34]. ViPTree analysis likewise showed no meaningful genomic similarity between Kane and any reference genome, and Kane did not consistently associate with a particular reference lineage across analyses, although it always fell among tailed dsDNA phages. vConTACT3 assigned Kane to the class *Caudoviricetes* but placed it in a predicted novel lineage distinct from that of Satoshi, Kolob, and Ushijima, with no assignment to a currently recognized order, family, subfamily, or genus.

Due to a lack of close identifiable relatives in genome-level comparisons, *Vibrio* phage Kane was more difficult to place taxonomically than the other three phages. Its highly divergent genome, distinct genome organization, and myovirus-like morphology distinguish it from the others characterized in this study. We therefore propose Kane as the sole currently known member of the new genus *Laievirus*, the name being derived from Lā‘ie, Hawai‘i, where Kane was isolated. A formal proposal to establish this genus and the single associated species, *Laievirus kane*, has been submitted to the ICTV.

### Genome Annotation and Gene Content

Satoshi and Kolob each contained 47 predicted coding DNA sequences (CDSs), Ushijima 49, and Kane 84, and none encoded tRNA genes (Table **1**). Named CDSs are listed in Supplementary Table 1. All predicted CDSs in the *Mokushimavirus* phages (Satoshi, Kolob, Ushijima, and CKB-S2) were encoded on a single strand, whereas Kane’s were divided between both strands (Table **1**).

The published CKB-S2 annotation contains 49 CDSs [15]. Our comparison revealed an additional boundary-spanning head-closure gene (nucleotides 32,637–34,713 and 1–134) with full-length homology to the head-closure proteins of Satoshi, Kolob, and Ushijima, bringing its total CDS count to 50.

Using the corrected CKB-S2 annotation, comparison of the predicted CDSs of the *Mokushimavirus* members revealed 52 protein families, seven of which were variable, lacking a homolog in at least one of the four *Mokushimavirus* phages. For the purposes of this paper, these seven families have been labeled F1–F7 (Table **2**). All seven were annotated as hypothetical proteins in each genome encoding them, with no biological functions yet identified.

**TABLE 2.** Protein families variable among the *Mokushimavirus* phages. Length is in amino acids (F5 is 129 aa in Ushijima and 131 aa in CKB-S2). A dash indicates that no homolog was predicted in that genome. Three families are unique to a single phage and have no homolog in the other three: F4 (Ushijima), F6 (CKB-S2), and F7 (CKB-S2). Satoshi and Kolob encode no unique protein.

| **Family** | **Length (aa)** | **Satoshi** | **Kolob** | **CKB-S2** | **Ushijima** |
|---|---|---|---|---|---|
| F1 | 152 | YGF25860.1 | YGF25904.1 | BAV81121.1 | — |
| F2 | 70 | YGF25862.1 | — | — | YGF26048.1 |
| F3 | 87 | — | YGF25902.1 | BAV81123.1 | YGF26047.1 |
| F4 | 118 | — | — | — | YGF26050.1 |
| F5 | 129–131 | — | — | BAV81119.1 | YGF26052.1 |
| F6 | 60 | — | — | BAV81115.1 | — |
| F7 | 82 | — | — | BAV81134.1 | — |

In Satoshi, Kolob, and Ushijima, F1–F5 were clustered within a 0.8–1.7 kb segment of each genome (Fig. **5**; coordinates in Supplementary Table 2B), downstream of a conserved GNAT-family N-acetyltransferase gene and interspersed among several other conserved genes (91–100% amino acid identity among the three phages), including a winged helix-turn-helix (wHTH) DNA-binding protein gene (accessions in Supplementary Table 2B). Immediately downstream of the wHTH gene, Satoshi and Kolob encoded F1, whereas Ushijima encoded F4; these families shared no detectable similarity by BLASTp, and neither had a recognizable remnant elsewhere in the other genome. F2 and F3 lay further downstream and likewise showed no detectable similarity, and F5 lay upstream of the wHTH gene in Ushijima only. Variation in gene content among Satoshi, Kolob, and Ushijima was thus concentrated in this single segment. CKB-S2 encoded F1, F3, and F5 (Table **2**) within the corresponding segment of its genome, whereas F6 lay a few genes upstream of the segment and F7 lay several genes downstream of it. The possible origins of the variable protein families are considered in the Discussion.

**FIG 5.**
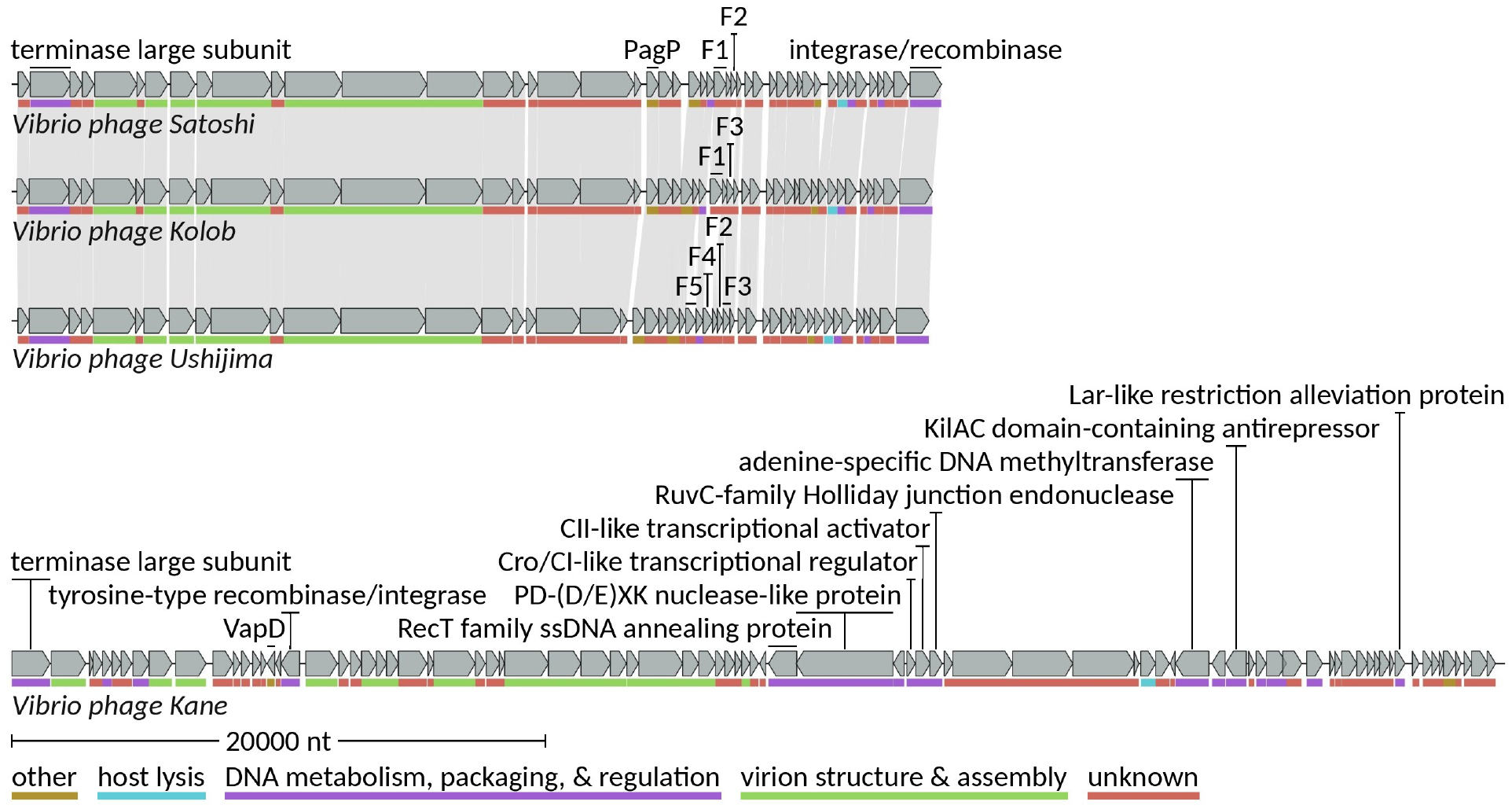
Comparative genome maps of *Vibrio* phages Satoshi, Kolob, Ushijima, and Kane, generated with LoVis4u. Gray arrows are predicted coding sequences, drawn in the direction of transcription. Colored bars beneath the arrows indicate functional categories (see key).

Comparative genome mapping with LoVis4u (Fig. **5**) revealed extensive gene-content conservation and synteny among Satoshi, Kolob, and Ushijima, whereas Kane exhibited a substantially different genome organization and lacked extensive synteny with the other three. Across all four novel phage genomes, genes with assigned functions were organized into distinct functional modules associated with either DNA metabolism, packaging, and regulation, or with virion structure and assembly (Fig. **5**).

Kane’s genome included several genes encoding proteins associated with integration and lysogeny-related regulation, including a tyrosine-type recombinase/integrase, a Cro/CI-like transcriptional regulator, a KilAC domain-containing antirepressor, and a CII-like transcriptional activator (Supplementary Table 1). Satoshi, Kolob, and Ushijima also encoded integrases/recombinases (Supplementary Table 1) and multiple DNA-binding proteins. Notably, all three encoded a conserved 83-aa protein (YGF25877.1, YGF25887.1, and YGF26032.1) with a lambda repressor-like helix-turn-helix fold (IPR010982), classified as a Cro/C1-type domain (IPR001387) in Satoshi and Kolob (Supplementary Table 3A), and HHpred matches included phage repressors such as P22 C2 (PDB 1ADR; Supplementary Table 3B), making it a candidate Cro/C1-like regulator.

The BACPHLIP algorithm [35] strongly predicted temperate lifestyles for all four novel phages (temperate scores of 0.91–0.99; Table **3**), consistent with the uniformly turbid plaque morphology and the integrase genes identified in each genome. In agreement with BACPHLIP, VIBRANT likewise classified all four genomes as lysogenic. The absence of tRNA genes from these genomes also lends support to this proposed lifestyle, as temperate phages tend to carry fewer tRNAs than virulent phages, whose codon usage diverges more from that of the host [36].

**TABLE 3.** BACPHLIP lifestyle predictions for the four phages. Scores are BACPHLIP random-forest outputs. The Satoshi genome was rotated to begin at nt 980 so that its integrase gene, which spans the ends of the deposited sequence, was contiguous for the analysis.

| Phage | Virulent Score | Temperate Score |
|---|---|---|
| Satoshi | 0.0375 | 0.9625 |
| Kolob | 0.0875 | 0.9125 |
| Kane | 0.0125 | 0.9875 |
| Ushijima | 0.05 | 0.95 |

BLASTn comparison of each phage genome against the complete genome of *V. coralliilyticus* OCN008 (CP048693–CP048695) [29] found no significant nucleotide similarity, arguing against induction of a resident OCN008 prophage as the source of any of the four phages.

Because temperate phages can alter host phenotypes through lysogenic conversion [25], the four genomes were examined for genes associated with bacterial virulence. Satoshi, Kolob, and Ushijima each encoded a homolog of PagP, a lipid A palmitoyltransferase associated with resistance to host antimicrobial peptides [37] (Supplementary Table 1). Alternatively, Kane encoded a homolog of VapD (YGF25980.1), another virulence-associated protein [38].

The PagP homologs of Satoshi and Kolob (identical in sequence) and of Ushijima shared 33.6–34.6% amino acid identity with PagP-annotated phage proteins, but their closest relative, a “hypothetical protein” of *Vibrio* phage CKB-S2 (BAV81113.1; 87.0–88.3% identity; Supplementary Table 2A), was not PagP-annotated. CKB-S2 therefore appears to encode a PagP homolog that was previously annotated as hypothetical.

The highest-scoring match to Kane’s VapD homolog was annotated only as “virulence factor” (*Enterobacter hormaechei*; 79.6% amino acid identity), and the highest-scoring match annotated explicitly as “Endoribonuclease VapD” was from *Neisseria bacilliformis* (59.6% identity; Supplementary Table 2A).

To test these annotations independently of BLASTp, the Satoshi/Kolob PagP homolog and the Kane VapD homolog were searched against experimentally determined structures (PDB70) using HHpred (Table **4**). The top match for the PagP-like protein was *Escherichia coli* PagP (PDB 3GP6); no other hit had *E <* 10*^−^*^5^. The top four matches for the Kane VapD-like protein were VapD structures from bacteria. Lower-ranked matches to Cas2-family endoribonucleases were consistent with a shared ferredoxin-like fold [39].

**TABLE 4.** HHpred matches of the Satoshi/Kolob PagP homolog and the Kane VapD homolog to experimentally determined structures. HHpred (MPI Bioinformatics Toolkit) searches against PDB70 (the Protein Data Bank clustered at 70% sequence identity), 26 Oct 2026. Identity is calculated over aligned positions. The Satoshi PagP homolog (YGF25854.1) is identical in amino acid sequence to Kolob’s and was not searched separately.

| PDB match | Source organism | Probability (%) | E-value | Query region aligned | Identity (%) |
|---|---|---|---|---|---|
| **Satoshi/Kolob PagP homolog** (146 aa; YGF25854.1/YGF25910.1) |  |  |  |  |  |
| 3GP6, PagP | *Escherichia coli* | 99.84 | 3.6 × 10<sup>−19</sup> | 30–146 | 14 |
| **Kane VapD homolog** (93 aa; YGF25980.1) |  |  |  |  |  |
| 6ZI1, VapD | *Haemophilus influenzae* | 99.97 | 7.4 × 10<sup>−30</sup> | 1–93 | 37 |
| 5DYR, VapD | *Xylella fastidiosa* | 99.96 | 5.7 × 10<sup>−28</sup> | 1–90 | 30 |
| 9KAN, VapD | *Campylobacter jejuni* | 99.96 | 4.9 × 10<sup>−28</sup> | 1–93 | 27 |
| 3UI3, VapD | *Helicobacter pylori* | 99.95 | 1.8 × 10<sup>−27</sup> | 1–93 | 35 |

Screening of the four genomes against the VFDB, CARD, and NCBI antimicrobial resistance gene databases using ABRicate (minimum 60% nucleotide identity and 50% coverage) detected no additional matches to known virulence, toxin, or antimicrobial-resistance genes.

Of Kane’s 84 CDSs, 49 (58.3%) had BLASTp matches (*E ≤* 10*^−^*^5^) to phage proteins (37, including 20 to *Vibrio* phages) or *Vibrio* bacterial proteins (21), with nine matching both (Supplementary Table 2A).

Of the 20 CDSs with matches to *Vibrio* phage proteins, only one, the hypothetical protein YGF25928.1, matched a protein from a *V. coralliilyticus* phage: Okinawan *Vibrio* phage CKB-S1 (42.7% amino acid identity; Supplementary Table 2A), which infects strain P1 [15] from the Great Barrier Reef, Australia [8]. The same protein also matched *Vibrio* phage vB_VpP_1, which is listed in GenBank as infecting *Vibrio parahaemolyticus*.

Among the nine CDSs with matches to both phage and *Vibrio* bacterial proteins were two adjacent, co-oriented genes that overlapped by 29 bp and encoded a PD-(D/E)XK nuclease-like protein (YGF25950.1) and a RecT family ssDNA annealing protein (YGF25951.1). The RecT family protein shared 40.3% amino acid identity with a RecT recombinase from *Vibrio harveyi*. The 1,207-residue nuclease-like protein had BLASTp matches only in its C-terminal region (residues 755–1207); the best matches with an assigned function were RecE-family exodeoxyribonucleases (exonuclease VIII) from enterobacteria (Supplementary Table 2A). These annotations and the adjacent arrangement of the two genes suggest a nuclease-annealase pair, reminiscent of the RecE-RecT system of the *E. coli* Rac prophage and the Exo-Red*β* system of *E. coli* bacteriophage *λ* [40], a hypothesis which is considered further in the Discussion.

Kane also encoded a predicted RuvC-family Holliday junction endonuclease, potentially involved in resolving Holliday junctions and other branched DNA intermediates formed during phage replication and recombination [41]. This shared 55.5% amino acid identity with a RuvC-family protein from *Photobacterium halotolerans* (Supplementary Table 2A).

Two other Kane proteins with predicted functions may help it counter host restriction-modification systems: an adenine-specific DNA methyltransferase and a Lar-like restriction alleviation protein (Supplementary Table 2A). Phage-encoded methyltransferases can protect phage DNA from cleavage by host restriction endonucleases by methylating the sequences these enzymes recognize [42]. Lar, a protein of the *E. coli* Rac prophage, instead acts on the host’s own restriction enzyme: it reduces cleavage of unmodified DNA by the Type I enzyme EcoKI and increases methylation of that DNA by the same enzyme [43]. If functional in *V. coralliilyticus*, the two proteins would be complementary, one shielding recognition sites in the phage DNA and the other dampening restriction activity in the host.

Beyond individual proteins, gene order was conserved in two pairs of adjacent Kane CDSs despite the absence of close genome-level relatives. In each pair, both Kane CDSs and the adjacent, co-oriented genes they matched in a reference *Vibrio* phage encoded hypothetical proteins (Table **5**). Pharokka additionally predicted tail-fiber functions for both proteins encoded by the second pair, but the annotation “hypothetical protein” was retained because none of their BLASTp matches had an assigned function. Stop-start overlaps and gaps are given in Table **5**. The possible significance of these arrangements is considered in the Discussion.

**TABLE 5.** Adjacent Kane CDS pairs with conserved gene order in reference *Vibrio* phages. Protein identity, query coverage, and E-values are from BLASTp. *Vibrio* phage 1.205.O._10N.222.51.A7 (MG592575) was isolated on *Vibrio tasmaniensis*, while *Vibrio* phage 1.052.A._10N.286.46.C3 (MG592433) was isolated on *Enterovibrio norvegicus* [30].

| **Kane protein** | **Reference protein** | **Amino acid identity (%)** | **Query coverage (%)** | **E-value** |
|---|---|---|---|---|
| **Pair 1**: *Vibrio* phage 1.205.O._10N.222.51.A7. Stop-start junction: 1-bp overlap in Kane (TAATG) and 1-bp gap in the reference (TAGCATG) |  |  |  |  |
| YGF25944.1 | NVP1205O_43 | 27.8 | 73.4 | 1.35 × 10<sup>−32</sup> |
| YGF25945.1 | NVP1205O_42 | 56.4 | 91.3 | 8.35 × 10<sup>−17</sup> |
| **Pair 2**: *Vibrio* phage 1.052.A._10N.286.46.C3. Stop-start junction: identical 4-bp ATGA overlap in Kane and in the reference |  |  |  |  |
| YGF25956.1 | NVP1052A_26 | 69.4 | 50.8 | 2.80 × 10<sup>−21</sup> |
| YGF25957.1 | NVP1052A_25 | 41.4 | 100 | 8.75 × 10<sup>−16</sup> |

## DISCUSSION

To the best of the authors’ knowledge, while a previous study [20] identified *V. coralliilyticus* phages from the Kona Coast of Hawai‘i Island, this is the first study to isolate *V. coralliilyticus* phages from O‘ahu. Phage isolation from this island is especially significant because, although *Montipora* white syndrome has been reported across the Hawaiian archipelago, it was noted to be “particularly prevalent in Kaneohe Bay, Oahu” [44]. Lā‘ie, the site from which the phages were isolated, lies only about 28 km from the bay.

All four phages produced plaques on *V. coralliilyticus* strain OCN008, a known etiological agent of *Montipora* white syndrome, but in preliminary tests none produced plaques on strain H1, which, like OCN008, was isolated from Kāne‘ohe Bay. This is consistent with the strain-level specificity reported for other *V. coralliilyticus* phages [20].

Along with being the first from O‘ahu, Satoshi, Kolob, Ushijima, and Kane are the first isolated *V. coralliilyticus* phages described as temperate. Notably, the Okinawan phage CKB-S2, which clusters within *Mokushimavirus*, was previously described as lytic on the basis of clear plaque morphology [15], yet encodes an integrase, a marker gene for temperate phages [23], like its Hawaiian relatives. Temperate phages can replicate productively or lysogenically depending on host physiology and phage density [23], so clear plaques under one set of assay conditions do not exclude a temperate lifestyle. CKB-S2 may therefore be capable of lysogeny, which would complicate its use in phage therapy, an application suggested by its discoverers [15], because temperate phages can increase host virulence and render lysogens immune to further phage infection [22].

In addition to similar lifestyles, members of the proposed genus *Mokushimavirus* would also be expected to have similar virion structures. Ushijima, the only Hawaiian member imaged, is podovirus-like (Fig. **1**A), and Satoshi and Kolob likely share this morphology: they encode the same virion proteins, including a predicted internal virion protein and tail spike, but lack the annotated tail sheath, tape measure, and baseplate proteins found in the myovirus-like Kane. CKB-S2, however, was described as myovirus-like by the study that isolated it, which reported a tail length of 105.88 nm from a single micrograph without reporting particle number or measurement variability [15]. The original report also contained an apparent discrepancy. Although the study described CKB-S2 as myovirus-like in morphology, its terminase phylogeny grouped it with the short-tailed *Vibrio* phage VP2, and the paper listed CKB-S2 as “Family *Podoviridae*?” in its classification table. The annotated CKB-S2 genome encodes the same internal virion protein as Satoshi, Kolob, and Ushijima (98% amino acid identity), and no tail sheath, tape measure, or baseplate proteins. These genomic features are far more consistent with a podovirus-like phage, but the discrepancy with the published micrograph and stated measurements cannot be resolved without re-imaging CKB-S2.

With the exception of a single 0.8–1.7 kb segment in which small hypothetical genes were gained, lost, or replaced, Satoshi, Kolob, and Ushijima were otherwise highly conserved in gene content, order, and orientation. The variable protein families F1–F5, which appear within the segment, illustrate the modular, mosaic architecture typical of phage genomes. In this architecture, genomes are assembled from discrete modules (single genes or groups of genes) that can vary independently from the rest of the genome. Each genome is therefore a mosaic: related phages share a common backbone of core genes but carry alternative modules (non-core genes) at some positions, and a single genome can contain regions with different evolutionary histories [45].

Mapping the variable protein families onto the proposed phylogeny (Figs. **3** and **4**) suggests that F1, F3, and F5 were each lost once, on separate branches, which is the most parsimonious explanation for their uneven distribution among the phages. F1 appears to have been ancestral to the four-phage clade and was lost in the Ushijima lineage, which instead encodes the unrelated gene F4 at the same position. F3 was lost in Satoshi, and F5 was lost in the lineage leading to Satoshi and Kolob. F2 (Table **2**) required at least two events, such as independent losses in Kolob and CKB-S2, or a gain in the ancestor of the three Hawaiian phages, followed by a loss in Kolob. F6 had no BLASTp matches outside CKB-S2 and most likely arose in the CKB-S2 lineage, whereas F7 shared amino acid identity with hypothetical proteins from at least two of the related *Vibrio* phages in Fig. **2** (1.138.O._10N.261.48.A1 and 284E43-1), consistent with its loss in the common ancestor of the Hawaiian lineage.

Replacement of F1 by the unrelated F4 at the same position is a plausible outcome of the mechanisms thought to generate phage mosaicism. One precedent comes from mycobacteriophages, in which mosaic modules are frequently single genes and recombination junctions lie at or near gene boundaries, a pattern attributed to illegitimate recombination followed by selection against recombinants that disrupt gene function [46]. The F1/F4 replacement matches this pattern: one single-gene module replaced another. A second precedent comes from *Pseudomonas aeruginosa* Mu-like phages, in which accessory ORFs are confined to a few shared loci, where different phages carry different ORFs presumably acquired by horizontal transfer [47]. The variable genes of Satoshi, Kolob, and Ushijima follow the same pattern: they are confined to a single segment, in which each phage carries its own combination of F1–F5. As is common for non-core genes [45], none of F1–F7 had a predicted function, so whether they benefit the phages that carry them remains to be determined.

Unlike the *Mokushimavirus* phages, Kane has no close relatives, yet 39 of its predicted proteins were assigned functional annotations based on homology to previously reported proteins, indicating distant relationships to other phages. For example, BLASTp of the major capsid protein reveals homology to the major capsid protein of *Salmonella* phage 118970 (NC_031940.1), in support of Kane’s myovirus-like morphology [48].

Like F1–F5 in the *Mokushimavirus* members, the two Kane gene pairs conserved in other *Vibrio* phages (Table **5**) are consistent with modular exchange between phages, although here the exchanged modules comprise two adjacent genes rather than one. Both pairs have spacing within the range associated with translational coupling [49], which may help explain why each pair was transferred as a unit. While vertical inheritance from a common ancestor cannot be excluded, horizontal transfer by recombination from unsampled phages seems the more likely explanation for the genes’ origins, given that Kane has no close relatives at the genome level.

Kane’s genome also encodes a candidate recombination system that could account for such transfers. Two adjacent genes encode a RecE-like nuclease and a RecT-family annealing protein, a pairing that resembles the RecE-RecT system of the Rac prophage and its functional counterpart in bacteriophage *λ*, Exo-Red*β*. In both systems, the nuclease degrades the 5*^′^* strand at double-stranded DNA ends to leave 3*^′^* single-stranded overhangs, which the annealing protein then pairs with complementary sequence [40]. Rad52-like recombinases such as RecT and Red*β* act independently of host RecA and promote gene shuffling among temperate phages [50]. They also recombine diverged sequences more efficiently than host machinery: in *λ*, the Red system recombined sequences diverged by 22% far more efficiently than the host RecABCD pathway [51]. Consistent with this, lambdoid phages that encode their own recombinases have more mosaic genomes, resulting from recent recombination events, and larger pan genomes than those that rely on host recombination functions [52]. Acquisition of a gene pair would require the incoming DNA to be joined to the recipient genome at its flanks, either by homeologous recombination between diverged flanking sequences or by illegitimate recombination at short or no homology followed by selection for functional recombinants [45, 51].

Kane further encodes a RuvC-family Holliday junction endonuclease, which could resolve the branched intermediates formed if an incoming segment were joined to the recipient genome by homologous or homeologous recombination at its flanks [41]. As a temperate phage, Kane’s genome is more likely to encounter other phages and prophages within the same host cell, which facilitates gene transfer by recombination [50]. Taken together, Kane’s predicted recombination functions, although assigned by homology, provide a plausible route for the acquisition of the gene pairs described above.

Kane encodes a fuller set of predicted lysogeny regulators than the other three phages. Its Cro/CI-like transcriptional regulator may participate in lytic-lysogenic switching, its CII-like transcriptional activator may promote entry into the lysogenic cycle, and its KilAC domain-containing antirepressor could promote the transition from lysogeny to the lytic cycle by counteracting repressor activity [53]. Satoshi, Kolob, and Ushijima lack the CII-like and antirepressor-like regulators found in Kane, but each encodes a candidate Cro/C1-like regulator, along with additional DNA-binding proteins of unassigned function. These additional DNA-binding proteins may be regulators too divergent to detect by homology, and the switch in these phages may involve regulatory mechanisms not represented in current databases.

Temperate phages, like those described in this study, can alter bacterial virulence through lysogenic conversion, the best-known example being cholera toxin, which is encoded by the filamentous phage CTX*ϕ* in *Vibrio cholerae* and causes the watery diarrhea thought to aid transmission [25]. Phages have also been implicated in the virulence of other *Vibrio* species: a myovirus-like phage increased the virulence of *Vibrio harveyi* strains toward prawn larvae [54]. In *V. coralliilyticus*, genomic analyses have identified prophages encoding toxin-like genes homologous to those of pathogenic *V. cholerae*, leading to the proposal that prophages contribute to virulence in this species [26].

Among the proteins predicted for Satoshi, Kolob, and Ushijima, the PagP-like proteins are of interest as candidate virulence factors that may influence the physiology and pathogenicity of the host bacterium *V. coralliilyticus* during lysogeny. Because CKB-S2 also encodes a PagP-like homolog, the gene appears to be conserved across all four members of *Mokushimavirus*.

PagP enzymes transfer palmitate from phospholipids to lipid A, the hydrophobic membrane-anchoring component of lipopolysaccharide (LPS), which is a major component of the outer membrane of Gram-negative bacteria, like *V. coralliilyticus* [55]. In experiments with *Salmonella enterica*, disruption of *pagP* increased outer-membrane permeability during exposure to a vertebrate cationic antimicrobial peptide (CAMP) of the innate immune system, indicating that lipid A palmitoylation impedes CAMP uptake across the outer membrane, thereby providing the bacteria with resistance to these peptides [37, 55].

This mechanism could be relevant to coral infection since coral species such as *Acropora digitifera* (*A. digitifera*) secrete antimicrobial peptides (AMPs) into their mucus to defend against bacterial invaders. Under low-salinity conditions, similar to those within coral mucus, the *A. digitifera*-encoded AMP, digitiferin, was found to exhibit bactericidal activity against *V. coralliilyticus* [6]. *M. capitata*, the coral species infected by OCN008, encodes a predicted digitiferin homolog [6]. PagP-mediated lipid A palmitoylation has also been proposed to provide an adaptive response to Mg^2+^ limitation [55]. If these phage-encoded proteins are expressed during lysogeny and function as lipid A palmitoyltransferases, they may increase the ability of *V. coralliilyticus* to resist coral immune defenses and withstand environmental stress.

Within the genome of Kane, a different candidate virulence factor was found: a VapD-like protein. VapD (Virulence-associated protein D) possesses ribonuclease activity, and its expression was previously linked to biofilm development in the bacterium *Xylella fastidiosa* [38]. If expressed during lysogeny, the Kane VapD protein could influence biofilm formation in *V. coralliilyticus*, and, in turn, pathogenicity. Of note, a VapD homolog has also previously been described in the *Vibrio harveyi* prophage phi345 [56].

Further research will be necessary to fully understand how these four phages interact with *V. coralliilyticus*. Of particular interest is whether the candidate virulence factors they carry, the PagP-like proteins of Satoshi, Kolob, and Ushijima and the VapD-like protein of Kane, enhance the pathogenicity of the bacterial host toward corals such as *M. capitata*. Establishing lysogens of OCN008 and comparing them with cured strains in assays of lipid A modification, biofilm formation, and susceptibility to coral antimicrobial peptides, followed by coral challenge experiments, would show whether these genes are functional and whether they influence disease outcome. By characterizing four temperate phages of *V. coralliilyticus* OCN008, a known agent of *Montipora* white syndrome, this study lays the groundwork for testing how viruses might shape coral disease.

## MATERIALS AND METHODS

### Bacterial Strains, Media, and Growth Conditions

Two strains of *V. coralliilyticus* were used in this study: OCN008, previously isolated from a *Porites compressa* coral fragment from Kāne‘ohe Bay, Hawai‘i [21], and H1, previously isolated from a *Montipora capitata* coral fragment, also from Kāne‘ohe Bay [57].

Strains were maintained by streaking on Zobell marine agar (Marine Agar 2216, HiMedia Laboratories, Mumbai, India) and were routinely cultivated in liquid marine broth (Marine Broth 2216, MilliporeSigma, Burlington, MA, United States) overnight at 30 *^◦^*C with shaking. Long-term stock cultures were stored at *−*80 *^◦^*C in marine broth supplemented with glycerol.

### Sample Collection

Environmental samples, comprising approximately 200 mL of sediment and 300 mL of seawater, were collected in coastal reef areas 15–20 m offshore of Temple Beach (21*^◦^*39*^′^*01*^′′^*N 157*^◦^*55*^′^*22*^′′^*W) in Lā‘ie, Hawai‘i, at a depth of 2–3 m. Sampling on 22 June 2024 yielded Satoshi and Kolob, and sampling on 20 February 2025 yielded Ushijima and Kane.

### Enrichment Method

Samples contained in 1-L jars were agitated on an orbital shaker at 210 rpm for 1 h at 20 *^◦^*C to suspend benthic bacteria and their associated bacteriophages into the overlying water column. Following agitation, a 50 mL aliquot of this seawater was clarified by centrifugation at 4,700 rpm for 15 min, and the supernatant was passed through a 0.22 µm pore-size membrane filter to remove larger particles and isolate the viral community, before being added to 10 mL of marine broth inoculated with 1 mL of *V. coralliilyticus*. This enrichment culture was gently agitated at 210 rpm and maintained at 20 *^◦^*C for 24 h to allow *V. coralliilyticus* phage replication.

After 24 h of incubation, the clarification (4,700 rpm, 15 min) and filtration (0.22 µm) steps were repeated using the enrichment culture instead of seawater. A 2 mL aliquot of the resulting filtrate was used to initiate a second round of enrichment, by inoculation into 40 mL of marine broth containing 1 mL of *V. coralliilyticus*. The second-round enrichment culture was subsequently clarified (4,700 rpm, 15 min) and filtered (0.22 µm).

### Phage Plaque Assays

Following the third round of filtration, double agar overlay plaque assays were performed for phage detection and purification. A 40 µL aliquot of filtrate was added to 300 µL of *V. coralliilyticus* broth culture and incubated for 20 min at 20 *^◦^*C to allow phage adsorption. A 15 µL aliquot of this mixture was then added to 3 mL of soft molten top agar containing 0.5% agarose and 3.5% marine broth powder supplemented with 1 mM MgCl_2_, and 1 mM CaCl_2_ after being pre-cooled to 45–50 *^◦^*C. The solution was mixed gently and poured over the surface of a marine agar plate. Top agar was allowed to solidify for 5 min before plates were moved to an incubator. Plates were incubated for 48–72 h at 26 *^◦^*C and then observed for plaque formation. A small agar plug containing a single, isolated plaque was aseptically transferred using a sterile pipette tip and suspended in 100 µL of phosphate-buffered saline (PBS). The suspension was serially diluted tenfold by transferring 10 µL into 90 µL of PBS at each step, after which 300 µL of *V. coralliilyticus* broth culture was added to each dilution and plaque assays were performed. Plaque purification via plaque assays was repeated at least three times to ensure a pure phage stock.

### Production of High Titer Phage Lysates and Titering

High-titer lysates were prepared by flooding confluent-lysis plates with 5 mL of PBS. Plates were swirled gently and incubated overnight at 4 *^◦^*C, followed by 1 h of incubation at 20 *^◦^*C with periodic swirling. The lysate was collected aseptically using a Luer-Lock syringe and filtered through a 0.22 µm syringe filter. Phage titers were determined using full-plate overlay plaque assays.

### DNA Extraction

Phage genomic DNA was extracted from high-titer lysates (*>*10^9^ PFU/mL) using the Norgen Biotek Phage DNA Isolation Kit (Norgen Biotek Corp., Thorold, ON, Canada) according to the manufacturer’s instructions. DNA concentration and purity were assessed using a NanoDrop 2000c spectrophotometer (Thermo Scientific).

### Next-Generation Sequencing and De Novo Assembly

Whole-genome sequencing of three of the four phages, designated Satoshi, Kolob, and Kane, was performed by SeqCenter (Pittsburgh, PA, USA). Sequencing libraries were prepared using the Illumina DNA Prep kit with custom IDT 10 bp unique dual indices and a target insert size of 280 bp. Samples were sequenced on an Illumina NovaSeq X Plus platform to generate 2*×*151 bp paired-end reads. Demultiplexing, quality control, and adapter trimming were performed using bcl-convert v4.2.4. All subsequent genome processing and analysis for these three phages were performed at BYU-Hawaii.

DNA from the fourth phage, designated Ushijima, was barcoded using the NEBNext Ultra II library prep kit (New England Biolabs), followed by pooling and paired end sequencing on the Illumina iSeq at BYU in Provo, Utah.

All read preprocessing and genome assembly steps for phages Satoshi, Kolob, and Kane were performed in Galaxy v26.0.1.dev1 [58]. Because sequencing depth greatly exceeded that required for phage genome assembly, raw sequencing reads were subsampled using seqtk sample v1.5 [59] with a random number generator seed of 4 and a subsampling fraction of 0.1 to reduce dataset size for downstream analyses. Reads were then filtered and trimmed using fastp v1.0.1 [60] with adapter auto-detection enabled. Filtering was performed using a qualified Phred score threshold of 20, an unqualified base percentage limit of 30%, and a minimum read length of 50 bp. Quality trimming was applied at the 5*^′^* end, 3*^′^* end, and by right-end sliding-window trimming, each using a window size of 4 and a mean quality threshold of 20. Automatic polyG tail trimming was enabled.

Filtered and trimmed reads were assembled de novo using SPAdes v4.2.0 [61, 62] with custom k-mer sizes of 77, 99, and 127. Assembly statistics were evaluated using QUAST v5.3.0 [63], and assembly graphs were visualized using Bandage-NG v2022.09 [64].

For phage Ushijima, raw fastq files were assembled using Geneious 2025 [65] at default de novo assembly settings.

### Genome Annotation and Analysis

Genome annotation was performed using DNA Master v5.23.6 [66] and GeneMarkS v4.28 [67], with BLASTp [68] used to manually assign putative gene functions. Functional assignments were further evaluated using Pharokka v1.3.2 [69] in Galaxy Europe (UseGalaxy.eu; Galaxy v26.0.1.dev1) [58]. The encoded amino acid sequences of select genes were also evaluated using HHpred [70] in the MPI Bioinformatics Toolkit to improve functional annotation. Protein sequences of interest were analyzed with InterProScan [71] through the EMBL-EBI web service (InterPro 110.0). Default settings were used for all annotation programs.

Phage genome sequences were queried against the NCBI Virus nucleotide database (nt_viruses) using BLAST+ [72], with the megablast task [73, 74] used for Satoshi, Kolob, and Ushijima and the discontiguous megablast task used for Kane.

Intergenomic similarities between the phage genomes and their top matches on BLASTn, ranked by total score, were calculated using the VIRIDIC web server (Moraru Phage Lab) [75] and visualized as a heatmap. Species and genus thresholds were set to 95% and 70%, respectively, following standard bacteriophage taxonomic practice [33].

Whole-genome proteomic tree analysis was performed using ViPTree v4.0 [76]. Complete genome sequences of the newly isolated phages and the top BLASTn-matched phage genomes for each isolate were uploaded in FASTA format and analyzed with the dsDNA prokaryotic virus reference set using default parameters. ViPTree generated a proteomic tree based on genome-wide sequence similarities computed with tBLASTx [76]. The resulting tree was used to assess the placement of newly isolated phages relative to reference phages.

Phylogenetic relationships between phages Satoshi, Kolob, Ushijima, and selected reference phages were further evaluated using terminase large subunit (TerL) amino acid sequences. Sequence alignment and all subsequent phylogenetic analyses were performed in MEGA12 [77]. TerL sequences were aligned using the MUSCLE algorithm [78]. Positions with less than 95% site coverage were removed using the partial-deletion option, leaving 504 positions in the final alignment. A maximum-likelihood phylogeny was inferred using the Jones-Taylor-Thornton amino acid substitution model [79]. Of two candidate trees generated using neighbor-joining [80] and maximum-parsimony methods, the tree with the higher log-likelihood was used as the initial tree for the heuristic search. Branch support was assessed using 1,000 bootstrap replicates [81].

A genome-wide gene-sharing network analysis was performed using vConTACT3 v3.2.4 [82] with the reference database v232 [83] within the Windows Subsystem for Linux environment to assess the taxonomic placement of the newly isolated phages relative to reference viruses.

Comparative genome maps were generated in the same environment using LoVis4u v0.2.0 [84]. Predicted proteins were manually assigned to functional categories for this visualization.

Phage lifestyle was predicted using BACPHLIP v0.9.6 [35], a random forest classifier based on conserved protein domains, and VIBRANT v1.2.1 [85]. The Satoshi genome was rotated to begin at nt 980 so that its integrase gene was contiguous. To test for related prophages, each phage genome was queried by BLASTn against the complete genome of *V. coralliilyticus* OCN008 (CP048693–CP048695) [29].

Genomes were screened for virulence, toxin, and antimicrobial-resistance genes using ABRicate v1.4.0 [86] against VFDB [87], CARD [88], and the NCBI AMRFinderPlus database [89], with minimum thresholds of 60% nucleotide identity and 50% coverage.

### Negative-Stain Transmission Electron Microscopy

Micrographs were obtained for representative *Vibrio* phages Ushijima and Kane. Three µL of each phage sample were directly applied to the carbon side of glow-discharged copper support C-flat holey carbon grids and allowed to adsorb for 1 min. Excess liquid was removed by side-blotting on Whatman No. 1 filter paper. Grids were then washed twice by gently dabbing the carbon side on two 20 µL droplets of 1*×* PBS successively, side blotting after each application. 3% uranyl acetate stain was then applied to the grids in the same fashion, including an additional step where the grid was left to float carbon side down on a third drop of stain for 1 minute followed by side-blotting to dryness.

Grids were stored in a vacuum desiccator for at least 30 minutes before being transferred to a glass desiccator for storage. Samples were imaged on the 120 kV Talos L120C TEM at the Yale Cryo-EM Resource core. Viral dimensions were measured using Fiji (ImageJ v2.16.0) [90].

## ACKNOWLEDGMENTS

The authors thank the many individuals at Brigham Young University-Hawaii who contributed in various ways to the laboratory work described in this study. In addition, the authors thank Natalie Olsen of Brigham Young University for checking the annotation of Kane and assembling the genome of phage Ushijima. The authors also thank Dr. Blake Ushijima of the University of North Carolina Wilmington for providing the bacterial strains used. An earlier version of this manuscript was posted as a preprint on bioRxiv: https://doi.org/10.64898/2026.09.21.753358

## DATA AVAILABILITY STATEMENT

The nucleotide sequence data generated in this study have been deposited in the NCBI GenBank repository (https://www.ncbi.nlm.nih.gov/genbank/) under accession numbers PZ716015 (*Vibrio* phage Satoshi), PZ716016 (*Vibrio* phage Kolob), PZ716017 (*Vibrio* phage Kane), and PZ716018 (*Vibrio* phage Ushijima).

